# CITE-seq analysis reveals human cytomegalovirus and diabetes-associated adaptive NK cell alterations in cardiovascular disease

**DOI:** 10.1101/2024.03.22.581997

**Authors:** Sujit Silas Armstrong, Daniel G. Chen, Sunil Kumar, James R. Heath, Matthew J. Feinstein, John R. Greenland, Daniel R. Calabrese, Lewis L. Lanier, Klaus Ley, Avishai Shemesh

**Affiliations:** La Jolla Institute for Immunology, La Jolla, CA; Institute of Systems Biology, University of Washington, Seattle, WA; Immunology Center of Georgia, Augusta University, Augusta, GA, USA; Division of Cardiology, Department of Medicine, Northwestern University Feinberg School of Medicine; Department of Medicine, University of California, San Francisco, CA; Parker Institute for Cancer Immunotherapy San Francisco, CA; Department of Microbiology and Immunology, University of California, San Francisco, CA; Immunology Center of Georgia, Medical College of Georgia at Augusta University, Augusta, GA

## Abstract

Coronary artery disease (CAD) is a leading cause of mortality worldwide with Diabetes and human cytomegalovirus (HCMV) infection as risk factors. CAD’s influence on human NK cells is not well characterized. CITE-seq analysis of a CAD cohort of 61 patients revealed distinctly higher NK cell *SPON2* expression and lower *IFNG* expression in severe CAD patients. Interestingly, HCMV^+^ patients displayed lower *SPON2* expression while diabetes status reversed the HCMV effect. Diabetes led to diminished adaptive FcεRIγ^−/low^ NK cell frequencies and was associated with a higher PBMC *IL15*/*TGFB* transcript ratio, while TGFB increased in severe CAD. *SPON2* expression corresponded to changes in conventional vs. adaptive NK cell frequencies, and *SPON2/IFNG* ratio decreased in inflamed plaque tissue with an increased adaptive NK cell gene signature and was increased in severe CAD patients. Our results indicate that the *SPON2*/*IFNG* ratio and adaptive NK cell gene signature associated with stenosis severity or inflammation in CAD.

## Introduction

Coronary artery disease (CAD) represents a third of all cardiovascular diseases, with pathophysiology associated with atherosclerotic plaque formation in the arteries that supply blood to the heart^1^. CAD severity is reported to increase with diabetes, likely through plaque remodeling^2^, but the mechanisms are not fully understood. Several other CAD risk factors are HCMV infection^3^, smoking, obesity, increased LDL and reduced HDL, and high blood pressure^1^. CAD immune responses are described as increased Th1 responses, associated with increased IFN-γ and chronic low-grade inflammation, macrophage infiltration in the neo-intima, foam cell formation, and decreased smooth muscle cell proliferation^4,5^.

NK cells are innate lymphocytes that regulate other immune, non-immune, virally infected, and transformed cells by cytokine secretion and direct cell lysis^6,7^. NK cells are CD3^−^CD56^+^CD16^−/+^ lymphocytes with two main subsets in blood: immature CD56^bright^CD16^−^ and mature CD56^dim^CD16^+^ cells. In CAD, reduced human peripheral blood NK cell numbers correspond to increased low-grade cardiac inflammation and heart failure^8^. Mouse NK cells were shown to limit cardiac inflammation and fibrosis by arresting eosino-phil Infiltration^9^. Reduced blood NK cell numbers are reported in SARS-CoV-2 infection and are associated with an increased mature adaptive NK cell immunophenotype and gene signature^10–13^. Human adaptive NK cell differentiation accurses with HCMV infection or reactivation^10^. Mature adaptive NK cells are characterized by NKG2A^−^NKG2C^high^ expression, yet adaptive NKG2C^negative^ NK cells were reported in NKG2C-deficient humans, indicating other mechanisms of differentiation such as due to CD16 stimulation^11,14,15^. Adaptive NK cells can be further defined by the expression or lack of expression of the adaptor protein FcεRIγ. Adaptive FcεRIγ^−^ NK cells (g^−^NK cells)^16^ exhibit reduced protein expression of the IL-2 receptor beta-chain (CD122, *IL2RB*), and high expression of IL-32, CCL5, GZMH, and LAG3^10,17,18^. In a pre-clinical study of coronary atherosclerosis, adaptive NKG2C^+^CD57^+^ NK cells were associated with lower plaque volume, suggesting a protective function^19^. Nonetheless, there is limited data regarding the human NK cell gene expression profile and subset differentiation in CAD and with respect to HCMV status and other risk factors^20,21^. Thus, CAD-specific impact on NK cells remains unknown.

Here, we analyzed CITE-seq data from a CAD cohort of 61 patients to deconvolve the influence of CAD, diabetes, HCMV status, and other risk factors on human NK cells. We found higher NK cell *SPON2* mRNA expression in severe CAD patients (CAD^high^) that negatively correlated with NK cell *IFNG* mRNA expression. HCMV^+^ CAD^high^ patients displayed lower *SPON2* expression, while diabetes opposed HCMV impact. NK cell cluster analysis revealed that diabetes status reduced adaptive FcεRIγ^−/low^ NK cell frequencies, which was linked to an increased PBMC *IL-15*/*TGFβ* mRNA ratio, while both cytokines deferentially regulated Spon-din-2 expression in adaptive and non-adaptive NK cells. Correspondingly, higher NK cell *SPON2* expression strongly correlated with reduced adaptive NK cell frequencies and increased conventional NK cell frequencies in CAD. Analysis of arteriosclerosis plaque tissue revealed an increased adaptive NK cell gene signature and lower *SPON2*/*IFNG* ratio in inflamed plaques, whereas the *SPON2*/*IFNG* ratio increased with CAD stenosis severity. Thus, alterations in a NK cell adaptive gene signature and *SPON2*/*IFNG* ratio in CAD are markers of CAD-associated inflammation and stenosis severity.

## Results

### Increased NK cell *SPON2* expression in severe CAD patients

To study the relationship between CAD and NK cells in humans, we analyzed PBMC CITE-seq data from a cohort of 61 patients diagnosed with a low or high CAD severity score (percent stenosis of each artery segment; CAD^low^: 0-6, or CAD^high^: x > 30, and as was previously defined: GSE190570)^22^. Thirty-one patients were diagnosed with T2DM (type 2 diabetes). Therefore, the cohort’s patients were grouped as: I) CAD-^low^diabetes^−^; n = 16, II) CAD^low^diabetes^+^; n = 13, III) CAD^high^diabetes^−^; n = 18, and IV) CAD^high^diabetes^+^; n = 14 (CAD/diabetes groups). No significant differences in age, sex, and BMI were detected between the patient groups (Supplementary Table 1). None of the patients reported having prior heart failure and all patients exhibited a normal creatinine range, indicating normal kidney function^23^, with diabetes patients expressed lower creatinine levels^24^. To identify blood NK cells, we used CITE-seq surface protein expression^25^ and identified 12 immune cell clusters (Figure 1A). Cluster 4 (> 10,000 cells) was characterized as CD56^+^, CD16^+/−^, CD3^−^, CD19^−^, CD20^−^, CD14^−^, CD123^−^, and CD4^−^ cells (Figure 1A). Furthermore, cluster 4 cells expressed high levels of NK cell-related genes; *NKG7* (Natural Killer cell granule protein 7), *PRF1* (perforin), *GZMB* (granzyme B), *KLRG1* (KLRG1), *IL2RB* (CD122), and *TBX21* (T-bet) (Figure 1B)^17,26,27^. As these peripheral blood cells expressed CD56 and CD16 proteins and perforin and *KLRG1* mRNA, we concluded these cells are NK cells^28^.

**Figure 1:**
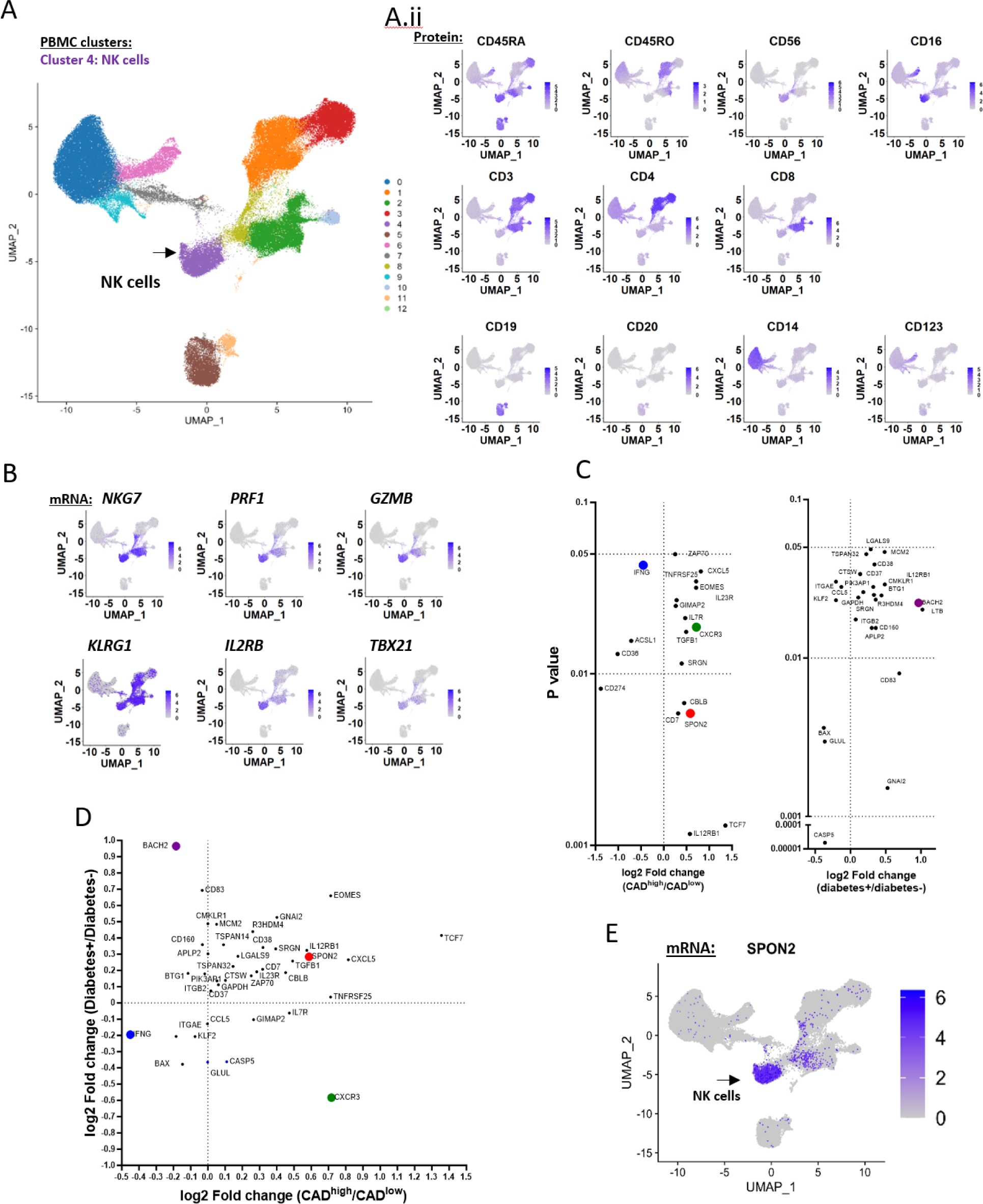
NK cell *SPON2* mRNA expression significantly increased in CAD. **A)** PBMC from 61 patients with or without diabetes or CAD were clustered by CITE-seq protein expression. **A.i)** uMAP of PBMC clusters based on CITE-seq protein expression. NK cells (purple cluster, black arrow). A.ii) uMAP of PBMC clusters for the specific markers CD45RA, CD45RO, CD56, CD16, CD3, CD4, CD8, CD19, CD20, CD14, or CD123. **B)** Single-cell RNA-seq analysis of the relevant NK cell-associated markers, *NKG7, PRF1, GZMB, KLRG1, IL12RB,* and *TBX21.* **C)** Dot plots displaying differential gene expression (DGE) analysis of patient’s NK cell mean gene expression between (left) diabetes^−^ vs. diabetes^+^ patients or (middle) CAD^low^ vs. CAD^high^ patients. X-axis: fold-change between patient groups, y-axis: p values (log10). Multiple t-tests, Mann Whitney test, compare ranks. **D)** Dot plot of CAD^high^/CAD^low^ fold change vs. diabetes^+^/diabetes^−^ fold change gene expression. *SPON2* (red), *IFNG* (blue), *CXCR3* (green), *BACH2* (purple). **E)** uMAP of *SPON2* mRNA expres-sion relative to PBMC clusters.

To identify CAD-specific variation in NK cell gene expression, we performed a differential gene expression analysis based on the patient’s NK cell mean gene expression between CAD^low^ vs. CAD^high^ (CAD^low^diabetes^−/+^ vs. CAD^high^diabetes ^−/+^) patients (Figure 1C). We discovered a significant increased expression of *IL12RB1*, *TCF7*, *CD7*, *SPON2*, *CBLB, SRGN*, *TGFB1*, *CXCR3*, *IL7R*, *IL23R*, *EOMES*, *GIMAP2*, *TNFR8F25*, *CXCL5*, and *ZAP70,* and a significant decrease in *IFNG, ACSL1, CD36, and CD274*^29^. As *SPON2*^30^ (red dot) and *CXCR3*^31^ (green dot) are reported in cardiovascular disease, and *IFNG* (blue dot) is reported with increased NK cell activation^32^, we focus on those genes. To check if the increased expression of these genes is associated with diabetes status, we compared diabetes^−^ vs. diabetes^+^ (diabetes^−^CAD^low/high^ vs. diabetes^+^CAD^low/high^) patients. In line with other publications, we found a high increase in *BACH2* (purple dot) expression in diabetes^+^ cases^33,34^ (Figure 1C, 1D). Yet, *SPON2*, *CXCR3*, and *IFNG* expression did not significantly change based on diabetes status alone.

We then checked if *SPON2* expression was specific to NK cells and found that *SPON2* transcripts were strongly expressed by the NK cell clusters with minimal expression by the T cell clusters (Figure 1E). Therefore, we concluded that increased CAD severity is associated with an increase in NK cell *SPON2 and CXCR3* expression, and lower *IFNG* expression.

### HCMV is associated with lower NK cell *SPON2* expression in CAD^high^diabetes^−^ but not in CAD^high^diabetes^+^ patients

To test if HCMV influences NK cell *SPON2* expression, we tested the patients’ anti-HCMV IgG1 serostatus. Of the 61 patients, 28 were HCMV seronegative (HCMV^−^), and 33 were HCMV seropositive (HCMV^+^). We identified a significantly higher number of HCMV^+^ cases in the CAD^high^diabetes^+^ group, [HCMV^−^ vs. HCMV^+^ cases: I) CAD^low^diabetes^−^ (8 vs. 8), II) CAD^low^diabetes^+^ (8 vs. 5), III) CAD^high^diabetes^−^ (10 vs. 9), and IV) CAD-^high^diabetes^+^ (2 vs. 12]) (Figure 2A.i). The increase in HCMV^+^ cases was age-independent and did not associate with high Systolic blood pressure (BP) levels or BMI score (Figure 2A.ii).

**Figure 2:**
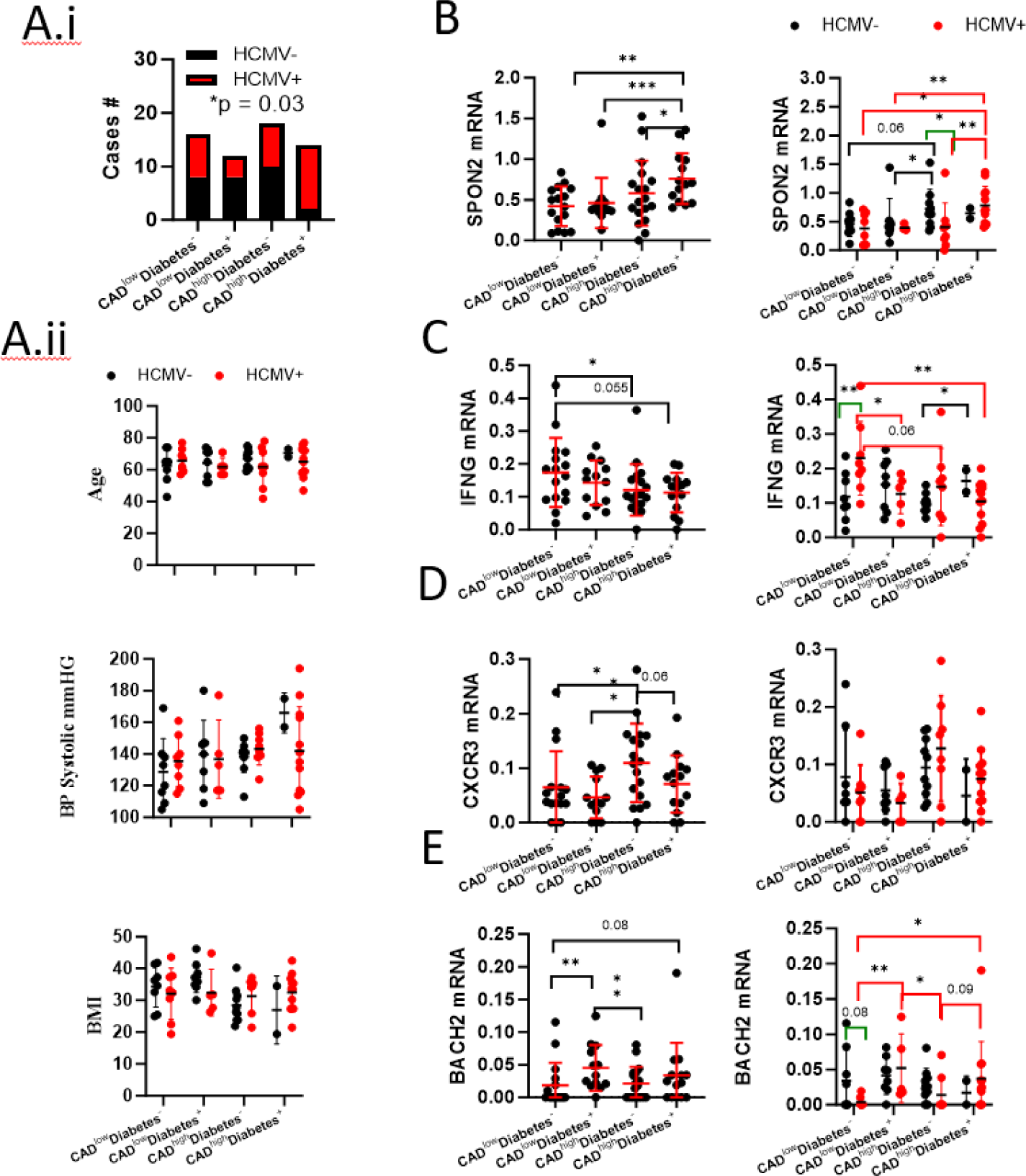
HCMV impact on NK cell *SPON2* mRNA expression. **A.i)**_Statistical analysis of patient’s HCMV serostatus (black: negative, red: positive) between the patients’ groups (Chi-square test). A.ii) Variation of age, BP systolic, or BMI (upper to lower) between the patients’ groups and HCMV serostatus. Variation of **B)** NK cell *SPON2*, **C)** NK cell *IFNG*, **D)** NK cell *CXCR3*, or **E)** NK cell *BACH2* mean expression per patient between the patients’ groups (left), and HCMV serostatus (right). Mean+/− S.D., Mann Whitney test, one-tail, *p < 0.05, ** p < 0.01, *** p < 0.001.

Plotting *SPON2* expression, based on the patients’ groups and HCMV serostatus, revealed lower *SPON2* expression in HCMV^+^ relative to HCMV^−^ CAD^high^ diabetes^−^ patients (Figure 2B), indicating that HCMV infection suppresses NK cell *SPON2* increase in severe CAD. In contrast, in CAD^high^diabetes^+^ patients, we did not reveal a decrease in NK cell *SPON2* expression in HCMV^+^ cases (Figure 2B), suggesting diabetes suppresses the influence of HCMV on NK cells *SPON2* expression during severe CAD. *SPON2* expression did not correlate to other risk factors such as age, blood pressure, or BMI (Figure S1A), indicating CAD-specific alterations that are impacted by HCMV and diabetes.

We then compared *IFNG*, *CXCR3*, and *BACH2* expressions between the patient groups and HCMV serostatus. *IFNG* levels were significantly elevated in HCMV^+^ relative to HCMV^−,^ CAD^low^diabetes^−^ patients, indicating that HCMV leads to NK cell activation (Figure 2C). Yet, diabetes or CAD attenuated *IFNG* expression independent of HCMV serostatus, suggesting a suppression of NK cell *in vivo* function or changes in NK cell subsets associated with reduced *IFNG* expression^32,35^. *CXCR3* expression increased in CAD^high^diabetes^−^ patients and exhibited reduced expression in CAD^high^diabetes^+^ patients, independently of HCMV serostatus (Figure 2D). *BACH2* expression was lower in HCMV^+^ relative to HCMV^−^ CAD^low^diabetes^−^ patients and increased in diabetes^+^ patients independently of HCMV serostatus (Figure 2E). Furthermore, *BACH2* negatively correlated to *IFNG* expression (Figure S1D). Thus suggesting diabetes status is associated with reduced NK cell terminal maturation or activation^10,33^. Accordingly, *IFNG* and *CXCR3* expression positively correlated to hsCRP (high-sensitivity C-reactive protein) levels, while *SPON2* expression negatively correlated with *IFNG* or *CXCR3* expression suggesting reduced inflammation^36^ (Figure S1B-D).

The results show that higher NK cell *SPON2* expression is CAD stenosis severity-specific marker, is impacted by HCMV serostatus, and is associated with reduced inflammation and NK cell maturation or activation. Further, it shows that diabetes opposes the negative impact of HCMV on *SPON2* expression in HCMV^+^ cases.

### HCMV-associated adaptive FcεRIγ^−/low^ NK cell frequencies are reduced in diabetes patients

To examine if our observations are associated with variations in the NK cell subpopulations, we clustered the NK cell population based on available CITE-seq panel protein expression We used markers previously reported with NK cell maturation, differentiation, and activation (e.g., CD56, CD16, CD25^10^, CD27^33,37^, CD2^14^, HLA-DR^11,15^). We identified five NK cell clusters (NK1-5) (Figure 3A). Gene expression analysis of NK cell maturation and differentiation-associated genes identified NK4 as immature NK cells based on the reduced expression of *FCGR3A* (CD16), *GZMA*, *GZMH*, *GZMB, BCL11B*, and *B3GAT1 (*CD57), with higher expression of *NCAM1* (CD56)^28^, *KLRC1* (NKG2A), *EOMES*, *CD27*, and *GZMK* relative to the other four clusters (Figure 3B)^14,17,18,37^. Accordingly, NK4 proportions significantly correlated with *GZMK* expression independently of HCMV serostatus (Figure S1E)^26^ and NK4 *CXCR3* expression decreased with diabetes^+^ status (Figure S1F). NK3 and NK5 displayed a mature conventional NK cell phenotype with reduced *FCERIG*, *BCL2*, *CD2*, *IL-32*, *and LAG3* expression^14,17,18^. NK5 differed from NK3 by reduced NCAM1 (CD56) expression, higher *GNLY* expression, and lower *BCL2* and *BCL11B* expression, and might represent less active cells^17,38^. NK1 and NK2 expressed higher levels of *CD2*, *IL-32*, *LAG3, GZMH, BCL2, and GNLY*^14,17,18^. NK2 expressed lower *IL2RB* (CD122) and *FCERIG* (FcεRIγ) levels and higher *LAG3* expression relative to other mature NK cell clusters, thus resembling mature adaptive FcεRIγ^−/low^ NK cells^10,39^. In line with the literature^10,39^, the frequencies of the NK2 cluster significantly increased in HCMV^+^ cases whereas other NK cell clusters did not show significant changes based on HCMV serostatus alone (Figure 3C).

**Figure 3:**
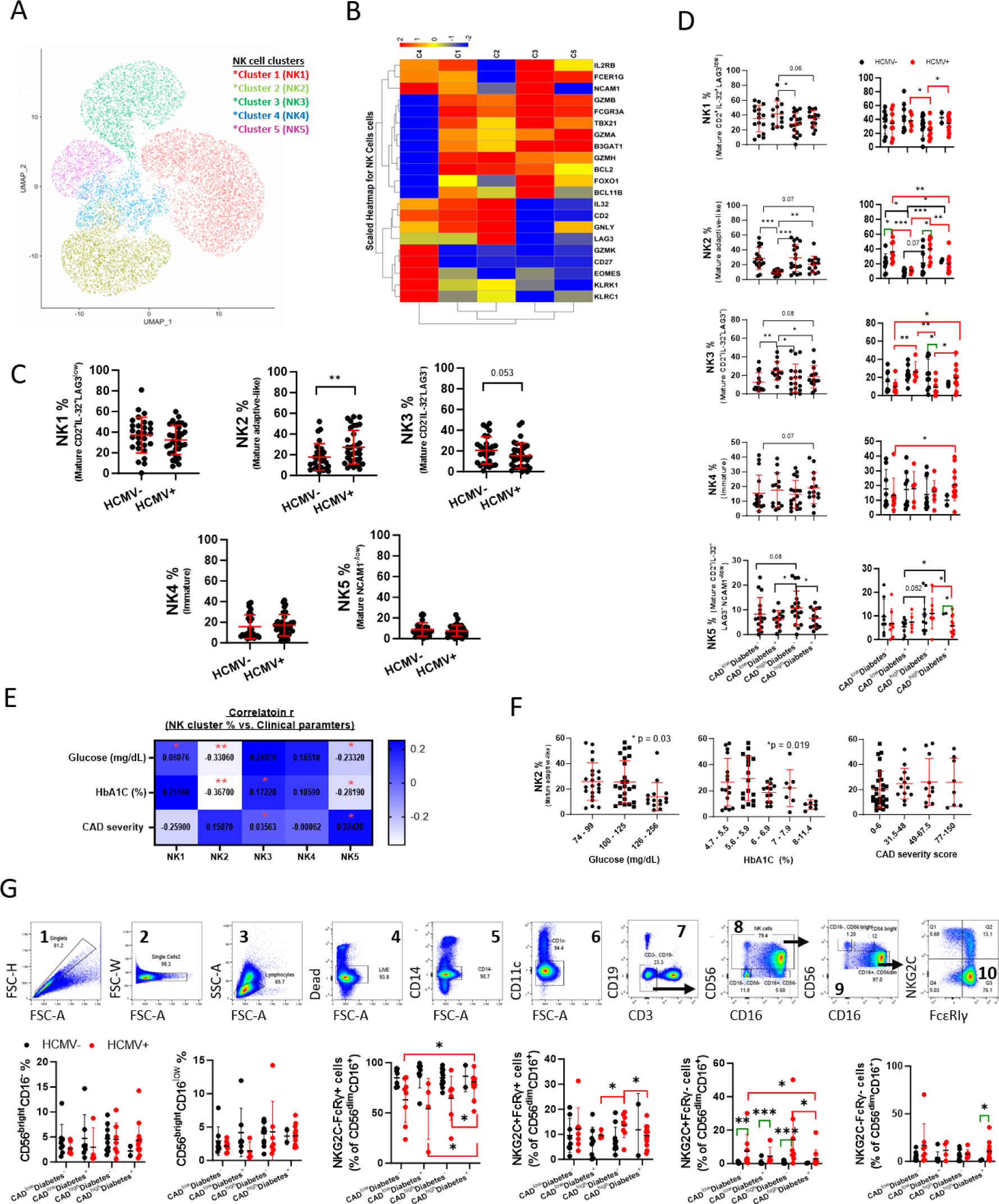
Variation in NK cell clusters associated with diabetes, CAD, and HCMV status. **A)** uMAP of NK cell clusters (Cluster 4 figure 1A) based on CITE-seq protein expression of CD56, CD16, CD25, CD27, CD2, or HLA-DR. **B)** heatmap of NK cell gene expression for the indicated identifier markers. **C)** Comparison of NK cell cluster proportions per patient (dot) between HCMV^−^ vs HCMV^+^ cases. **D)** Comparison of NK cell cluster proportions per patient (dot) between the patients’ group (left) and HCMV serostatus (right, black: HCMV^−^, red: HCMV^+^. **E)** Heatmap showing person correlation r values between NK cell clusters (C1-C5) frequencies and glucose (mg/mL), HbA1c %, or CAD severity score. **F)** Comparison of NK cell cluster 2 (NK2) proportions per patient (dot) between patients’ group based on (left to right): glucose (mg/mL), HbA1c %, or CAD severity score (one-way ANOVA). (* p <0.05). **G)** Cytek analysis of CAD cohort patients’ PBMC (n = 61, without CAD or diabetes status. **Upper panels: Cytek** gating strategy of PBMC: PBMC were gated to exclude doublets by FSC-A vs. FSC-H [1] and FSC-A vs. FSC-W [2], following by generating a lymphocyte gate [3] and removing of dead cells [4], CD14^+^ cells [5], and CD1c^+^ cells [6]. The remaining cells were then plotted by CD3 (T cells) vs. CD19 (B cells) [7], and CD3^−^CD19^−^ cells were plotted by CD56 vs. CD16 to identify NK cells [8]. NK cells were then gated as immature CD56^bright^CD16^−^, immature CD56^bright^ CD16^low^, and ma-ture CD56^dim^CD16^+^ [9]. Mature NK cells were plotted by FcɛRIγ vs. NKG2C to identify adaptive NK cell sub-sets [10]. **Lower panels:** Comparison of NK cell subsets percentages per sample between (upper panels) CAD and diabetes status, (middle panels) CAD^low^ vs. CAD^high^, or (lower panels) diabetes^−^ vs. diabetes^+^ pa-tients, and HCMV serostatus (black: negative, red: positive). Statistical analysis of panels C, D, and F, Mean+/− S.D, Mann, Whitney test, one-tail (* p <0.05, ** p <0.01, ***p <0.001, green bars: HCMV^−^ vs. HCMV^+^, Black bars: between HCMV^−^ patients’ groups, red bars: between HCMV^+^ patients’ groups).

We then compared NK cell cluster proportions between the patient groups and HCMV status (Figure 3D). NK2 (adaptive FcεRIγ^−/low^ NK cells) proportions significantly decreased in diabetes^+^ patients, independently of HCMV serostatus, indicating that diabetes status suppresses adaptive FcεRIγ^−/low^ NK cell frequencies. In contrast, NK3 (mature conventional *CD2*^−^, *IL-32^−^, LAG3*^−^ NK cells) proportions significantly increased in diabetes^+^ patients independently of HCMV serostatus. In line, diabetes^+^ status led to a reduced adaptive NK cell gene signature (Figure S1G) (e.g., higher *FCERIG* and *IL2RB* and lower *LAG3, IL32, CCL5,* and *GZMH* expression) while CAD^high^ status attenuated diabetes-associated variations. Accordingly, NK2 cluster frequencies negatively correlated and decreased with increased HbA1c or glucose levels (Figure 3E, 3F).

To validate our observation regarding adaptive FcεRIγ^−/low^ NK cell frequencies, we used Cytek flow cytom-etry to assess NKG2C^high^FcεRIγ^+^ or NKG2C^high^FcεRIγ^−^ NK cells frequencies in the CAD cohort patient’s PBMC samples (Figure 3G). Gating NK cells (CD3^−^CD19^−^CD56^dim^CD16^+^) based on NKG2C and FcεRIγ protein expression^10^ confirmed the reduced percentages of mature adaptive NKG2C^high^FcεRIγ^−/low^ NK cells in HCMV^+^diabetes^+^ patients (Figure 3G). Thus, we concluded that HCMV-associated adaptive FcɛRIγ^−/low^ NK cell frequencies are reduced in diabetes while severe CAD attenuates the impact of diabetes on NK cells.

### IL-15 and TGFβ regulate Spondin-2 expression in primary NK cells

Spondin-2 (Mindin), a secreted extracellular matrix protein encoded by the *SPON2* gene^40,41^. We have reported that mature adaptive NKG2C^high^ FcεRIγ^−^ and FcεRIγ^low^ NK cells express lower surface IL-2Rβ protein levels relative to mature non-adaptive NKG2C^−^FcɛRIγ^High^ and immature CD56^bright^CD16^−^ NK cells. Additionally, we reported that IL-2/15 receptor stimulation leads to FcεRIγ upregulation, which is inhibited by rapamycin (mTOR inhibitor) and TGFβ^10^. In line, NK2 (adaptive FcεRIγ^−/low^) cluster expressed lower *IL2RB* transcripts (Figure 3B). To determine if Spondin-2 expression is impacted by differential IL-2/15 receptor stimulation, we stimulated purified NK cells from adaptive NK cell-positive donors with increasing levels of IL-2 or IL-15 and measured intracellular Spondin-2 expression (Figure 4A). Note that the IL-2/15-dependent NK92 cell line expressed Spondin-2 (Figure S2A), yet short (1 day) IL-2, IL-15, or IL-12 stimulation of primary NK cells, which led to IFNγ expression, did not lead to Spondin-2 expression (Figure S2B), which required prolonged (3 day) IL-2/15 receptor stimulation (Figure S2C). Therefore, we compared Spondin-2 expression between immature (CD56^bright^ CD16^−^), mature (CD56^dim^ CD16^+^) non-adaptive NKG2C^−^ FcɛRIγ^High^ and adaptive NKG2C^high^ FcεRIγ^low^ or NKG2C^high^ FcεRIγ^−^ NK cells on day 3 (Figure 4A). Either IL-2 or IL-15 led to Sponidn-2 upregulation in a concentration-dependent manner, indicating regulation downstream of the IL-2/15 receptor. Adaptive NKG2C^high^ FcεRIγ^−^ or FcεRIγ^low^ expressed lower Spondin-2 protein relative to other NK cell subsets, thus showing differential Spondin-2 expression in NK cell subsets, which explains the lower NK cell *SPON2* expression in HCMV^+^ CAD^high^ diabetes^−^ cases relative to HCMV^−^ patients (Figure 2B).

**Figure 4:**
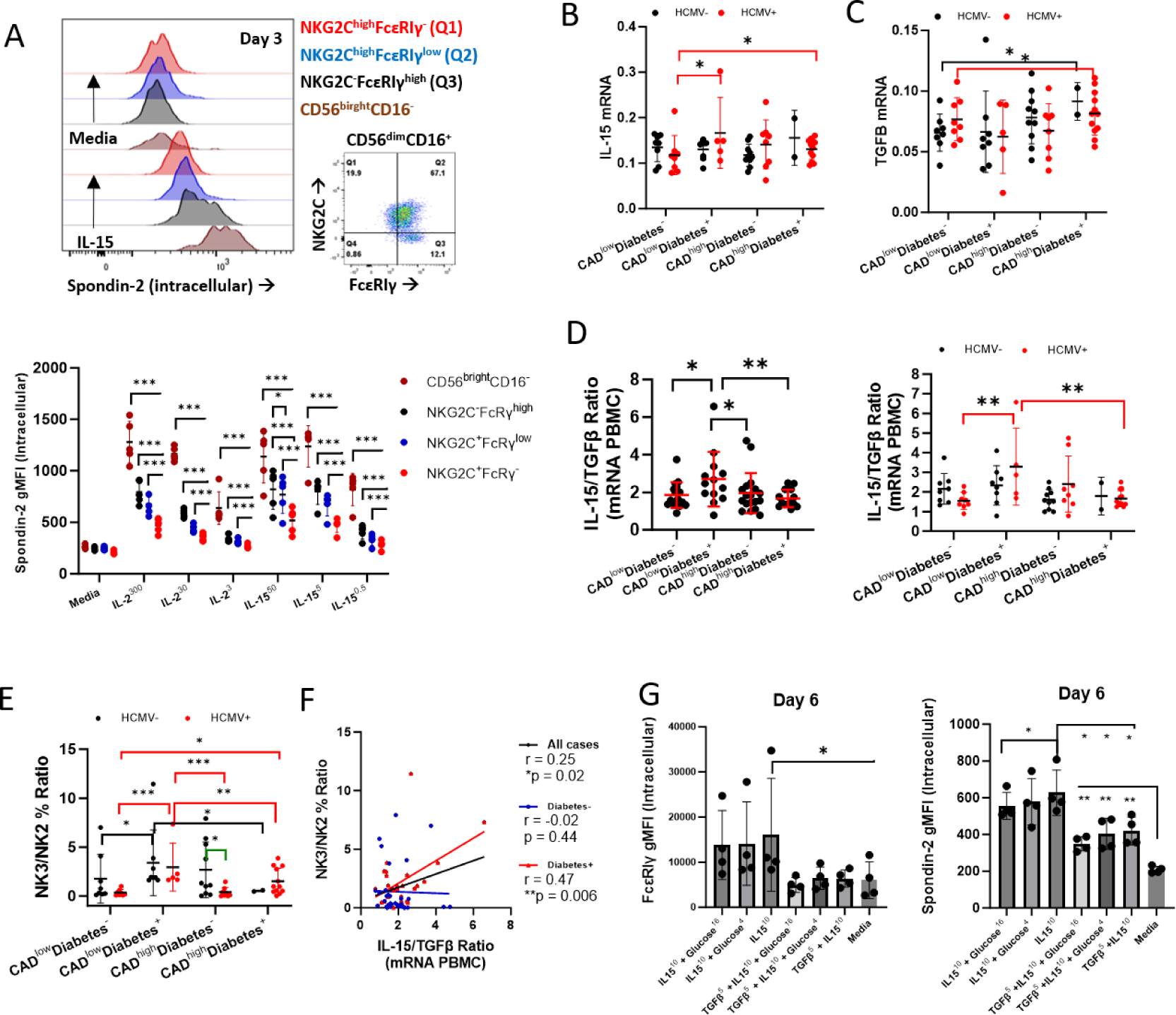
regulation of Spondin-2 expression by IL-15 and TGFβ in primary NK cells. **A)** Histograms of Spondin-2 intracellular expression between the defined NK cell subsets (color-coded) after 3 days of stimulation with IL-15 (50 ng/ml) or media without cytokines. The right dot plot represents the gating of mature CD56^dim^CD16^+^NK cells to identify adaptive NK cell subsets by NKG2C vs. FcεRIγ protein expression. Lower panel: IL-2 or IL-15 concentration-dependent expression of Spondin-2 between the defined NK cell subsets IL-2 (300, 30, and 3 U/ml), IL-15 (50 and 0.5 ng/ml) after 3 days of stimulation and relative to media without cytokines. Expression of CAD cohort’s PBMC **B)** IL-15 or **C)** TGFβ mRNA expression between the patients’ group and HCMV serostatus. **D)** PBMC IL-15/ TGFβ ratio between the patients’ group (left) and HCMV serostatus (left). **E)** NK3/NK2 cluster proportion ratio between the patients’ group and HCMV serostatus. **F)** Person correlation between IL-15/ TGFβ ratio and NK3/NK2 ratio in all cases (black), diabetes^−^ cases (blue), or diabetes^+^ cases (red). Upregulation of **G)** FcɛRIγ or Spondin-2 expression in purified primary NK cells stimulated for 6 days with IL-15 (10 ng/ml), with or without glucose (16 or 4 g/L) or TGFβ (5 ng/ml) relative to media without cytokines. Statistical analysis: panels B, C, D, and E: Mean+/− S.D, Mann, Whitney test, one-tail (* p <0.05, ** p <0.01, ***p <0.001, green bars: HCMV^−^ vs. HCMV^+^, Black bars: between HCMV^−^ patients’ groups, red bars: between HCMV^+^ patients’ groups).

We then examined the expressions of IL-2, IL-15, and TGFβ in the CAD cohort data. As we could not assess protein concentration in the plasma, which is subject to cytokine uptake, we assessed mRNA expression in PBMC. *IL-2* mRNA expression displayed no significant variations (data not shown*). IL-15* mRNA increased in CAD^low^diabetes^+^ and CAD^high^diabetes^+^, with HCMV serostatus dependency (Figure 4B), and positively correlated to glucose and HbA1c levels (Figure S2D). *TGFβ* mRNA increased in CAD^high^ cases with higher expression in CAD^high^diabetes^+^ patients (Figure 4C). Plotting the ratio of *IL-15* vs. *TGFβ* transcripts revealed a significant increase in CAD^low^diabetes^+^, but not in CAD^high^diabetes^+^ patients (Figure 4D). To determine if *IL-15*/ *TGFβ* ratio is associated with NK cell cluster variations in diabetes^+^ patients, we examined the ratio between NK3 vs. NK2 clusters relative to the patients’ groups. In line with the increase in *IL-15*/*TGFβ* ratio, we detected a significant increase in the NK3/NK2 ratio in CAD^low^diabetes^+^ cases, and a HCMV-associated increase in the CAD^high^diabetes^+^ groups relative to CAD^low^diabetes^−^ group (Figure 4E). Further, the *IL-15*/*TGFβ* ratio showed a significant positive correlation in the NK3/NK2 ratio in diabetes^+^ patients (Figure 4F). Thus, variations in the *IL-15*/*TGFβ* ratio explain the reduced proportions of the NK2 cluster in CAD-^low^diabetes^+^ patients and the attenuated decrease in CAD^high^diabetes^+^ patients and explain the opposing influence on NK cells between CAD or diabetes patient groups.

To further assess the regulation of Spondin-2 expression, we stimulated purified NK cells with IL-15 (10 ng/ml), with or without TGFβ (5 ng/ml) and in the presence of higher glucose concentrations (16 or 4 g/L relative to culture media) and measured Spondin-2 upregulation or FcεRIγ upregulation as a control at day 6. In line with our prior study^10^, TGFβ completely suppressed FcεRIγ upregulation by IL-15, while the increased glucose levels showed a negative influence on FcεRIγ upregulation (Figure 4G). In contrast, TGFβ partly suppressed Spondin-2 upregulation (Figure 4G), suggesting a differential regulation by TGFβ. Therefore, we examined Spondin-2 upregulation during mTOR inhibition by rapamycin (RAPA) and co-inhibition of FOXO1 (Figure S3E). In line with our previous findings ^10^, FcεRIγ upregulation was suppressed by rapamycin and was salvaged by FOXO1 co-inhibition. In contrast, rapamycin suppressed Spondin-2 upregulation whereas FOXO1 co-inhibition did not salvage its expression. Thus, showing differential regulation of Spondin-2 relative to FcεRIγ, which can explain the increase in NK cell *SPON2* expression in CAD^high^ patients in the presence of increasing TGFβ mRNA expression.

### *NK cell SPON2/IFNG ratio is* decreased in carotid plaque tissue and increased with CAD disease burden

As we discovered differential Spondin-2 upregulation in non-adaptive vs. adaptive NK cells and diabetes-associated alterations in the proportions of NK cell clusters, we tested whether changes in *SPON2* expression was associated with a particular NK cell subset^26,27^. *SPON2* expression negatively correlated with decreased NK2 (adaptive FcεRIγ^−/low^) proportions and positively correlated with increased NK3 (conventional) proportions (Figure 5A.i,ii). Thus, showing that alterations in *SPON2* mRNA expression in the CAD cohort are associated with changes in blood adaptive FcεRIγ^−/low^ NK cell frequencies.

**Figure 5:**
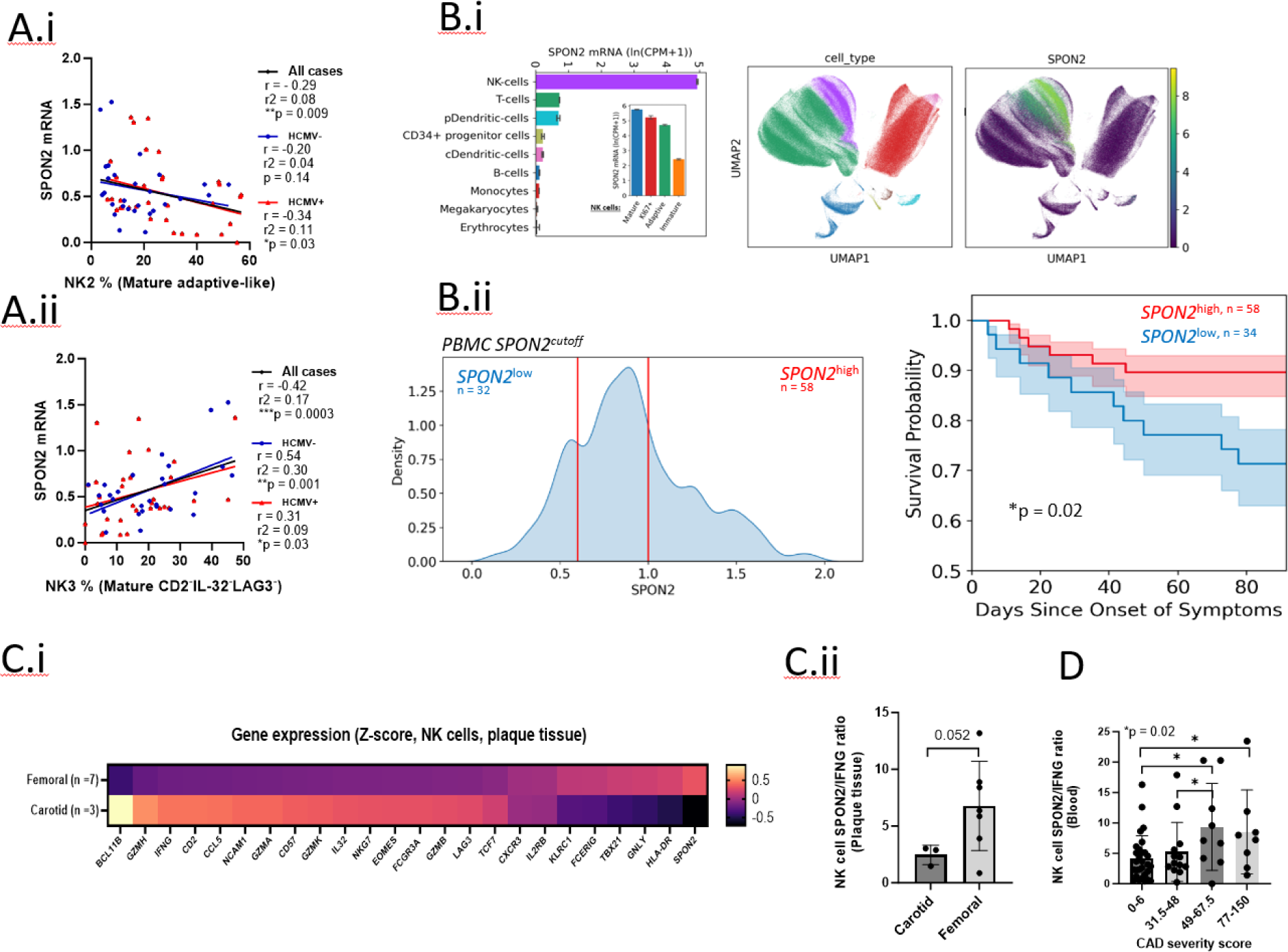
CAD NK cell *SPON2* mRNA expression in human NK cell clusters. **A)** Pearson correlation (one-tail) between patients’ mature adaptive NK2 (Upper) or mature conventional NK3 (lower) proportions vs. *SPON2* mRNA expression in 61 patients (black), HCMV^+^ patients (red), or HCMV^−^ patients (blue). **B)** PBMCs from COVID-19 patients (n = 216) were analyzed by single-cell RNA-seq. B.i; left: *SPON2* mRNA expression levels in NK cells relative to other defined cells, or in different NK cell subsets (inner panel), uMAP of PBMC clusters based on RNA expression (middle panel), and *SPON2* mRNA expression in the PBMC clusters (right panel). B:ii; Kaplan-Meier plot of survival probability based on *SPON2* high vs. low cutoff (right. SPON2-low; n = 32, *SPON2*-high; n = 58). **C)** NK cell gene expression between the carotid plaque (n =3) relative to the femoral plaque (n =7). Gene expression is displayed as mean z-score. C.ii: NK cell *SPON2* vs NK cell *IFNG* ratio between carotid (n = 3) and femoral (n= 7) atherosclerotic plaques. **D)** NK cell SPON2/IFNG mRNA ratio relative to CAD severity score. Statistical analysis: Mean+/− S.D, Mann Whitney test, one-tail, or person correlation (one-tail).

To further examine this observation, we examined *SPON2* expression in NK cell subsets during SARS-CoV-2 infection, reported to increase adaptive FcεRIγ^−/low^ NK cell frequencies in association with COVID-19 patient’s death, disease severity, and increased inflammation^10,42^. Analysis of single-cell RNA sequencing data revealed high *SPON2* expression in NK cells relative to other cell types during SARS-CoV-2 infection, while immature and adaptive NK cells expressed lower *SPON2* mRNA (Figure 5B.i). Furthermore, higher PBMC *SPON2* expression was significantly associated with COVID-19 patients’ survival (Figure 5B.ii, Supplementary Table 2). Analysis of NK cell *SPON2* expression relative to COVID-19 disease severity (WOS score: H = healthy, 1-2 = mild, 3-4 = moderate, 5-7 = severe), at different time points (T1 = diagnosis, T2 = follow-up, one week after diagnosis, and T3 = long-term follow-up, 2-3 months after initial diagnosis and should represent any final outcomes of disease), revealed a decreased *SPON2* expression in mild cases at T1 and T2, and a significantly lower expression in severe cases at T3, relative to healthy controls (H). In contrast, NK cell *IFNG* expression increased with disease severity at T1, T2, and T3 (Figure S3A.i). Analysis of NK cell *SPON2* expression at T3 relative to patient death revealed reduced *SPON2* expression in patients who died while *IFNG* expression increased (Figure S3A.iI). Thus, *SPON2* expression is lower in adaptive NK cells, higher *SPON2* expression is associated with better patient survival, and NK cell *SPON2* expression opposes NK cell *IFNG* expression. Thus, we concluded that higher NK cell *SPON2* expression might be associated with reduced inflammation linked to reduced NK cell activation and a lower adaptive NK cell gene signature.

To further examine our observations relevance to Atherosclerosis, we examined *SPON2* expression in the atherosclerotic plaque tissue (Figure 5D). Analysis of NK single-cell RNA sequencing data (GSE23407)^43^ of human carotid plaques (“Inflamed”, n = 3) relative to the femoral plaques (“Stable”, n = 7) revealed decreased *SPON2* mRNA expression in carotid plaque, while *BCL11b*, *B3GAT1 (*CD57), *IFNG,* and adaptive NK cell-associated gene (e.g. *GZMH*, *IL32*, *CCL5*, and *LAG3*) mRNA expression increased (Figure 5C.i). *IFNG* expression positively correlated with adaptive NK cell-associated genes (*LAG3*, *GZMH*, *CCL5*, *IL32)* and neg-atively correlated to *FCERIG* and *KLRC1 (*NKG2A) expression (Figure S3B.ii), whereas *SPON2* showed a negative correlation to *IL32* (n =10), and *CXCR3* expression exhibited a positive correlation to *KLRC1* (NKG2A), *HLA-DR*, *IL2RB* (CD122), and *FCER1G* and a negative correlation to *GZMH*, *LAG3*, and *BCL11B*. Thus, indicating that NK cell *SPON2* expression is not associated with an adaptive NK cell gene signature or “inflamed” atherosclerotic plaques. Plotting NK cell *SPON2* expression relative to NK cell *IFNG* expression revealed a decreased ratio in carotid plaques relative to femoral plaques, indicating that a higher *SPON2*/*IFNG* ratio is associated with less inflamed and more stable plaques (Figure 5D.ii). Thus, the data validates that higher NK cell *SPON2* expression is a marker of more stable plaques associated with increased stenosis^44^ while lower SPON2/IFNG ratio is associated with thin inflamed plaques.

We then assessed the NK cell *SPON2*/*IFNG* ratio in the CAD cohort patients based on CAD disease severity (combined percent stenosis of each artery segment score: I; 0-6 [n = 29], II; 30-48 [n = 13], III; 49-67.5 [n = 9, IV; 77-150 [n = 8]). The NK cell *SPON2*/*IFNG* ratio significantly increased in patients with higher disease burden (III: 49-67.5 and IV: 77-150, relative to I: 0-6 and II: 30-48). This indicates that the increased CAD stenosis is associated with a higher NK cell *SPON2*/*IFNG* ratio and reduced NK cell activation and reflect plaque accumulation and decreased inflammation.

## Discussion

Here, we analyzed CITE-seq data from PBMC collected from CAD patients with different disease severity (low vs. high) and with or without diabetes to study the impact on human NK cells by CAD and CAD-associated risk factors. NK cells are strongly influenced by HCMV infection, which can lead to the accumulation of adaptive NK cell subsets. Thus, we included HCMV serostatus in our analysis to characterize CAD-specific changes in NK cells or the impact of HCMV on NK cells during CAD.

We found that in CAD^high^ patients NK cell *SPON2* expression increased while *IFNG* mRNA decreased. Additionally, NK cell *SPON2* expression was suppressed in CAD^high^HCMV^+^ patients, while diabetes opposed the HCMV effect. Accordingly, diabetes led to reduced frequencies of adaptive FcɛRIγ^−^ NK cells, while CAD^high^ attenuated the diabetes impact. We then studied Spondin-2 protein (encoded by *SPON2* gene) upregulation in human primary NK cells for the first time and showed that IL-2/15 receptor stimulation led to up-regulation of Spondin-2, while adaptive NK cells, reported to exhibit reduced IL-2 and IL-15 sensitivity^10,45^, expressed lower Spondin-2. Furthermore, we found that PBMCs’ *IL-15*/*TGFβ* mRNA ratio corresponded to variations in the conventional/adaptive NK cell (NK3/NK2) ratio, and that Spondin-2 upregulation was differently suppressed by TGFβ, relative to FcɛRIγ upregulation. Moreover, *SPON2* negatively correlated to adaptive FcɛRIγ^−^ NK cell (NK2) frequencies while positively correlating with mature conventional NK cell (NK3) frequencies. Thus, showing that variation in sensitivity to IL-2 or IL-15 in NK cells impacts gene expression across NK cell subsets. We have recently reported that adaptive NK cells express lower surface levels of IL-2 receptor beta chain and exhibit lower mTOR activity^10^. IL-2 or IL-15 stimulation leads to FcεRIγ upregulation, which is inhibited by rapamycin or TGFβ, and shows a positive correlation to cell proliferation^10,46^. Thus, NK cell studies in association to human diseases such as CAD are required to address changes in IL-15 or TGFβ, as well as HCMV serostatus and adaptive NK cell subsets, to better understand disease-associated changes in NK cells.

The accumulation of adaptive NK cells and an adaptive NK cell gene signature is reported during severe COVID-19 and is associated with reduced blood NK cell numbers, increased inflammation, and patient death. Blood NK cell numbers are reported to decrease with low-grade cardiac inflammation^47^, while restored circulating NK numbers are associated with reduced cardiac inflammation^8^. NK cell numbers were also reported to decrease with inflamed carotid plaques relative to stable femoral plaques^43^. In a pre-clinical study of coronary atherosclerosis, adaptive NKG2C^+^CD57^+^ NK cells were associated with lower plaque volume, suggesting a protective function^19^. Interestingly, we found that the adaptive NK cell gene signature increased in the carotid, more inflamed, plaque tissue. Carotid plaques are thin (lower volume) relative to femoral plaques (higher volume), which are more stable^48^. Thus suggesting that the accumulation of adaptive NK cells might be associated with increased inflammation and a higher risk of plaque rupture^49,50^.

We found that during COVID-19 adaptive NK cells expressed lower *SPON2* transcripts. Accordingly, high PBMCs’ *SPON2* expression was associated with better patient survival, and NK cell *SPON2* levels decreased with patients’ death while *IFNG* increased. Our analysis revealed the same trend of increased *IFNG* and adaptive NK cell-associated gene expression in inflamed carotid plaques relative to stable femoral plaques. In the CAD cohort analysis, NK cell *IFNG* expression positively correlated to hsCRP levels, while *SPON2* showed a negative correlation to *IFNG*. Interestingly, the NK cell *SPON2*/*IFNG* mRNA ratio increases with CAD severity. In our study, CAD severity was assessed by the presence of stenosis in each artery segment and reflected the overall angiographic disease burden. Stenosis reflects the narrowing of blood vessels due to plaque burden^44^. Indeed, we also detected an increase in PBMC TGFβ mRNA in severe CAD cases. TGFβ is a potent pro-fibrotic and anti-inflammatory agent in CAD, and it increases with CAD severity^51^. Aortic stenosis is associated with increased TGFβ and fibrosis^52,53^. Thus, the NK cell *SPON2*/*IFNG* ratio might be an indicator of TGFβ impact during CAD.

Recently, several publications identified *SPON2* expression in human NK cells in scleroderma, melanoma, acute myeloid leukemia, and tuberculosis^26,27,54,55^. However, the clinical interpretation of NK cell *SPON2* expression was not defined. In humans, Spondin-2 plasma levels increase with major cardiovascular events risk^30^. Yet, Spondin-2 is downregulated in humans with failing hearts^56^ and *Spon2* knockout in mice increases cardiac risk ^56–58^. The discrepancy between the increase in Sponidn-2 in human plasma with CAD severity and mice might be explained through a dysregulated protective mechanism. In line, the extracellular matrix proteoglycan, Lumican, is reported to increase in cardiovascular patients, while Lumican knockout in mice increased mortality post-aortic banding and led to decreased *Spon2* expression^59^. We have shown here that in CAD, NK cell *SPON*2 expression is an indicator of stenosis severity and is reduced in inflamed plaques. Thus, human NK cell *SPON2* might play an important role in CAD protection and might be an indicator of reduced inflammation in other diseases, as we shown in COVID-19. Yet further research is required to validate if NK cell SPON2 has a direct impact in these conditions or is it only a bio-marker reflecting inflammatory status.

Overall, our results reveal the CAD-specific impact on human NK cells and the co-impact of CAD risk factors, such as diabetes, and HCMV infection (and the interplay between them) on CAD’s influence and show that an adaptive NK cell gene signature or NK cell *SPON2* expression are indicators of increase inflammation or stenosis severity, respectively.

## Acknowledgments and Sources of Funding

Studies were supported by the NIH HL P01 HL136275 and R35 HL145241 to K. Ley, the Parker Institute for Cancer Immunotherapy to A. Shemesh, the Joel D. Cooper Award from the International Society for Heart and Lung Transplantation to D.R. Calabrese, the Cystic Fibrosis Foundation Harry Schwachman Career Development Award CALABR19Q0 to D.R. Calabrese, the Veterans Affairs Office of Research and Development CX002011 to J R. Greenland, and National Heart, Lung, and Blood Institute HL151552 to J.R. Greenland.

## Author contributions

Conceptualization: A. Shemesh, L.L. Lanier, and K. Ley. Data curation: A. Shemesh, D. G. Chen, and S. S. Armstrong. Formal analysis: A. Shemesh, D. G. Chen, and S. S. Armstrong. Funding acquisition: L.L. Lanier, A. Shemesh, K. Ley, D.R. Calabrese, J.R. Greenland, and J.R. Heath. Investigation: A. Shemesh, D. G. Chen, S. S. Armstrong, and S. Kumar. Methodology: A. Shemesh, S. S. Armstrong, and D. G. Chen. Project administration: A. Shemesh L.L. Lanier, and K. Ley. Resources: K. Ley, M.F. Feinstein, and J.R. Heath. Supervision and validation: A. Shemesh, L.L. Lanier, and K. Ley. Visualization: A. Shemesh, S. S. Armstrong, and D. G. Chen. Writing, original draft: A. Shemesh. Writing, review & editing: A. Shemesh, S. S. Armstrong, D. G. Chen, D.R. Calabrese, J.R. Heath, J.R. Greenland, and K. Ley, L.L. Lanier. Supervision: A. Shemesh, L.L. Lanier, and K. Ley.

## Methods

### Sample Collection and Quantitative Coronary Angiography (QCA) Quantification

As was previously described, individuals between 40 and 80 years old suspected of having coronary artery disease were recruited from the Coronary Assessment in Virginia cohort (CAVA) through the Cardiac Catheterization Laboratory at the University of Virginia Health System, Charlottesville, VA, USA. Written informed consent was obtained from all participants, and the study received approval from the Human Institutional Review Board (IRB No. 15328). Peripheral blood samples were collected from these participants before catheterization. Patients underwent standard cardiac catheterization with specific views of the coronary arteries. QCA was performed using automatic edge detection to analyze various parameters related to stenosis, including minimum lumen diameter, reference diameter, percentage diameter stenosis, and stenosis length. Analysis was carried out by experienced investigators who were blinded to the study. The severity score was determined based on the percentage stenosis of each artery segment, and the scores were combined to determine the overall angiographic disease burden. Patients were classified as CAD high if their score was >30 and CAD low if their score was <6. Diabetes status was evaluated by hemoglobin A1c (HbA1C) percentage and blood glucose (mg/dL) levels. Blood samples were collected before the SARS-CoV-2 outbreak.

### Preparation of PBMC Samples

Peripheral blood samples were collected from coronary artery disease patients and individuals who underwent cardiac catheterization to exclude CAD. PBMCs were isolated from the blood samples using Ficoll-Paque PLUS (GE Healthcare Biosciences AB, Uppsala, Sweden) gradient centrifugation. Cell viability was assessed using Trypan blue staining, and the PBMCs were cryopreserved in a freezing solution (90% fetal bovine serum with 10% DMSO). Prior to analysis, the frozen PBMCs were thawed, and the viability and cell count were determined. Next, the tubes containing the PBMCs were centrifuged at 400 × g for 5 minutes. The cells were then resuspended in a combination of 51 AbSeq antibodies, with each antibody added at a volume of 2 μL and 20 μL of BD’s Stain Buffer solution^22^. This resuspension process was performed on ice for 30-60 minutes according to the manufacturer’s recommendations. Afterward, the cells were washed and counted once again. Out of the total 65 samples examined, 61 samples successfully passed the quality control assessment with a cell viability rate exceeding 80%. For each subject, the cells were tagged using a Sample Multiplexing Kit from BD. Biosciences. The kit included oligonucleotide cell labeling. The tagged cells were subsequently washed three times, mixed, counted, stained with the relevant antibody mix, washed three more times, and finally loaded into Rhapsody nano-well plates. Each plate accommodated four samples.

### Library Preparation and Single-cell RNA-sequencing

Pre-sequencing quality control (QC) was conducted using Agilent TapeStation high-sensitivity D1000 screentape. Each tube was cleaned using AMPure XP beads and was washed with 80% ethanol. The cDNA was then eluted, and a second Tapestation QC was performed, followed by dilution as necessary. The samples were combined into a pool and subjected to sequencing according to the recommended parameters: AbSeq with 40,000 reads per cell, mRNA with 20,000 reads per cell, and sample tags with 600 reads per cell. The sequencing was performed on an Illumina NovaSeq using S1 and S2 100 cycle kits (Illumina, San Diego, CA, USA) with specific dimensions (67 × 8 × 50 bp).

The resulting FASTA and FASTQ files were uploaded to the Seven Bridges Genomics pipeline (https://www.sevenbridges.com/apps-and-pipelines/, accessed on 9 November 2020), where data filtering was applied to generate matrices and CSV files. This analysis yielded draft transcriptomes and surface phenotypes of 213,515 cells involving 496 genes and 51 antibodies^22^. After removing cell doublets based on sample tags and undetermined cells, 175,628 cells remained. Further doublets were eliminated using Doublet Finder (https://github.com/chris-mcginnis-ucsf/DoubletFinder, accessed on 7 December 2020), resulting in 162,454 remaining cells. Additionally, 291 NK cells were excluded as they appeared to be doublets with myeloid cells. NK cells were defined based on the presence of CD56+, CD16-/+, and the absence of CD19-, CD3-, CD19-, CD4-, CD14-, and CD123-protein expression. 10494 NK cells (6.46% of total PBMCs) were successfully identified.

### Thresholding and Clustering

Antibody thresholds were determined for each antibody by assessing its signal in negative cells or deconvoluting overlapping normal distributions of the known major cell types. The function normalmixEM from the mixtools R package was used to deconvolve the overlapping distributions. Ridgeline plots were used to set the best threshold for each antibody. Thresholding helps remove noise from non-specific antibody binding and enables characterizing cells with the right cell surface phenotype. Before clustering, the data was batch-corrected using the Harmony (v0.1.1) package. The dimensionality reduction of UMAP (Uniform Manifold Approximation and Projection) was used to project the cells onto a 2D space. The UMAP algorithm was applied to the first four principal components obtained from Harmony. The minimum dis-tant parameter was set at 1. The Louvain clustering algorithm was used to cluster the cells based on their surface phenotypes. For NK cell clusters, cells were clustered based on the expressions of CD56, CD16, CD25, CD2, CD27, and HLA-DR. The resolution parameter was set to 0.08, and the random seed was set to 42 for the reproducibility of results. Five distinct populations were identified after clustering.

### Single-cell RNA-seq data analysis

RNA and ADT quantifications for the identified NK cells were analyzed in R using Seurat (v4.3.0). Antibody data were CLR normalized and converted to the log2 scale, while transcripts were normalized based on total UMIs and converted to the log2 scale. Feature plots were generated using Seurat’s FeaturePlot func-tion. Differential expression or correlation analysis was performed on patients’ mean gene expression values to calculate the fold-change expression or correlation between the defined groups, P-values were calculated using patients’ mean gene expression values between the defined groups and tested by the Mann-Whitney test (two-tails or one-tail as indicated in the figure legends), and plots were created using GraphPad 10. Heatmap was generated using the heatmap (v1.0.12) R package. Gene expression data were scaled across all samples. Genes that were expressed in less than 40% of patients were removed to avoid misinterpretation of the results. Atherosclerotic plaque data were obtained from GSE23407.^43^

### HCMV ELISA

Anti-cytomegalovirus IgG1 serostatus was assessed using the human Anti-cytomegalovirus IgG1 ELISA kit (CMV, Abcam, AB108724). Patients’ serum samples were diluted at 1:10 and tested according to the company’s protocol.

### Cytek analysis

Frozen PBMC samples from the same 61 patients with or without CAD or diabetes were used to analyze NK cell subsets by a 5 laser Cytek Aurora. LIVE/DEAD™ Fixable Blue Dead Cell Stain Kit (Invitrogen, cat. No: L34962) was used to exclude dead cells. AF647-conjugated anti-CD14 (BioLegend cat. No: 302046) and BV711-conjugated anti-CD1c (BioLegend cat. No: 331536) were used to exclude myeloid cells. BUV805-conjugated anti-CD3 (BioLegend cat. No: 612895) was used to exclude T cells, and PE/Fire 700-conjugated anti-CD19 (BioLegend cat. No: 302276) was used to exclude B cells. CD3-CD19-cells were gated using BV570-conjugatyed anti-CD56 (BioLegend cat. No: 362540) vs. BV785-conjugated anti-CD16 (BioLegend cat. No: 302046), and CD56+CD16-/+ lymphocytes were defined as NK cells as described in the figure legend. Percp-cy5.5-conjugated anti-NKG2A (BioLegend cat. No: 375126), Pacific-blue-conjugated anti-CD57 (BioLegend cat. No: 359608), PE-conjugated anti-NKG2C (BioLegend cat. No: 375004) and FITE-conjugated anti-FcεRIγ (intracellular, Millipore Sigma cat. No: FCABS400F), and BUV615-conjugated anti-NKG2D BD™ Biosciences cat. No: 751232). Surface staining and intracellular staining were done as previously described^10^.

### Primary NK cell culture and Spondin-2 protein expression

Human primary NK cells were isolated from Plateletpheresis leukoreduction filters (Vitalant, https://vi-talant.org/Home.aspx) by using the negative selection “RosetteSep Human NK Cell Enrichment Cocktail” kit (#15065; STEMCELL Technologies). Primary NK cells were cultured in CellGenix® GMP stem-cell growth media (SCGM, 20802-0500) supplemented with 1% L-glutamine,1% penicillin and streptomycin, 1% sodium pyruvate, 1% non-essential amino acids, 10 mM HEPES, and 10% human serum (heat-inactivated, sterile-filtered, male AB plasma; Sigma-Aldrich). Adaptive NK cell-positive donors were identified by analysis of NKG2C (FAB138P or FAB138A antibodies, R&D Systems) and FcεRIγ (FCABS400F antibody, Millipore)^10^ protein expression on live (Zombie red, 77475; BioLegend), CD3-negative (300318 antibody; BioLegend), CD56^dim^ (318322 antibody; BioLegend) CD16^+^ (302038 antibody; BioLegend) cells. NK cells were cultured for the indicated amount of time with or without human IL-2 (TECINTM; teceleukin, ROCHE), human IL-15 (247-IL/CF; R&D Systems), human TGFβ1 (580706; BioLegend), human IL-12 (219-IL;R&D Systems), D-(+)-Glucose (Millipore-Sigma, G7021), mTORC1 inhibitor (Calbiochem; rapamycin, 553210, IC50 = 0.1 µM), or FOXO1 inhibitor Calbiochem (AS1842856, 344355, IC50 = 33 nM). Surface or intracellular staining was done as previously described^32^. Spondin-2 (Mindin Antibody (A-10) sc-166868 PE), IFNγ (502516 antibody; BioLegend). NK92 cell culture or primary NK cells anti-CD16 beads stimulation was done as previously described^32^. Human TruStain FcX™ (Fc Receptor Blocking Solution, 422302, BioLegend) was used at 1:100 dilution for blocking nonspecific binding. Data acquisition was performed using an LSR-II flow cytometer and analysis was performed by using FlowJo.v10.

### COVID-19 patients’ analysis

The lifelines package (Davidson-Pilon, 2021) was used to plot Kaplan-Meier (KM) curves for patient survival probability. The date of death was measured as days after the onset of initial COVID-19 symptoms. KM curves were plotted for up to three months to display all dead patients. Patients were substituted for those whose comorbidity was known and further split into those with and without given comorbidities. These two separate groups of patients were utilized to compute KM curves. Statistics for survival analysis were determined using a chi-squared test as implemented via scipy.stats.chi2_contingency. Single-cell transcrip-tome-based UMAP projections were obtained from the study done by Su *et al*.^42^. Cell-type level analyses were done using the pre-labeled cell types in the dataset. Pre-labeled NK cells were sequestered for further analyses (IRB No. 20170658).

### Statistical analysis

GraphPad Prism 10 or R statistical programming were used to calculate statistical differences using a Mann Whitney test, one-tail, or Pearson correlation, One-way ANOVA test, or chi-square test (one-tail) as described in the figure legends (*p ≤ 0.05; **p < 0.01; ***p< 0.001). Data values represent patient mean gene expression. Unless otherwise indicated, all graphs show mean +/− population standard deviation (S.D). Patient groups’ data points were integrated into one graph to allow better visualization of relative changes between HCMV^−^ and HCMV^+^ patients. A Mann Whitney test comparing variables between patient groups was done on HCMV^−^ patients (CAD/diabetes groups: number of tests = 6) or HCMV^+^ patients (CAD/diabetes groups: number of tests = 6) or between HCMV^−^ vs. HCMV^+^ patients within each defined patient group (CAD/diabetes groups: number of tests = 4).

### Data availability statement

All CAD and Atherosclerosis related data are available at GEO: GSE190570 and GSE23407. COVID-19 data are available as published^42^.

## Supplementary tables, figures and figure legends

**Supplementary Table 1:**
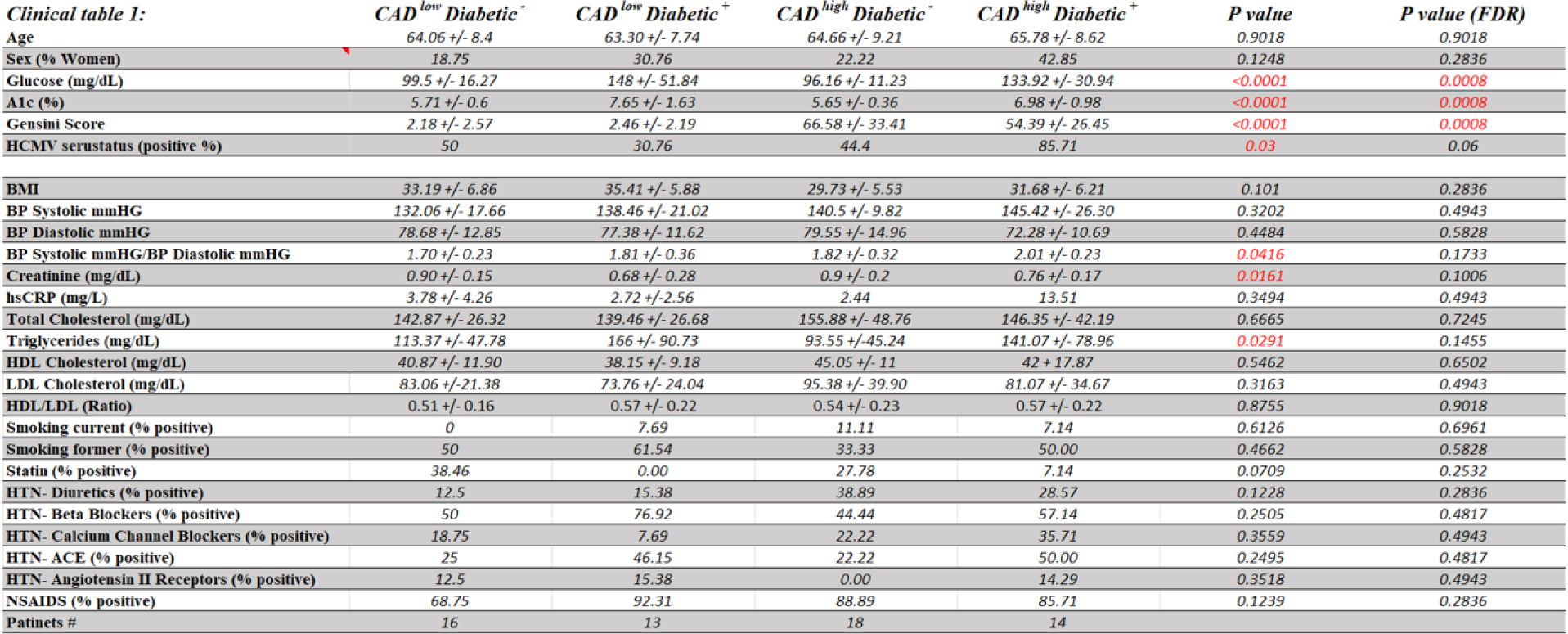
Clinical data of the CAD cohort’s CAD/diabetes groups based on CAD and diabetes status. Data are presented as mean +/− S.D or percentage of the population. Statistical analysis was done using the ANOVA-one-way or chi-square tests. p >0.05 was considered significant (red). Both the p-value and p-value after FDR correction are presented.

**Supplementary Table 2:**
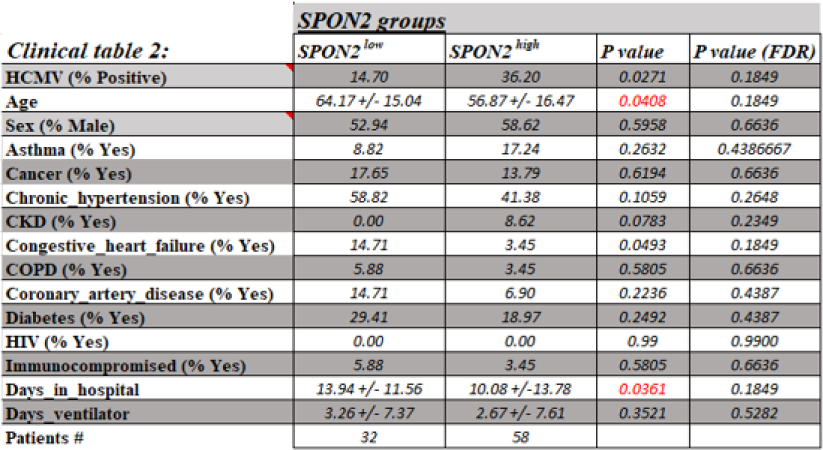
Clinical data of the COVID-19 cohort’s CAD, diabetes, or *SPON2* groups. Data are presented as mean +/− S.D or percentage of the population. Statistical analysis was done using the unpaired t-test or chi-square tests. p <0.05 was considered significant (red). Both the p-value and p-value after FDR correction are presented. Variations in patient numbers between the CAD or diabetes groups are due to a lack of diabetes status information.

**Supplementary Figure 1:**
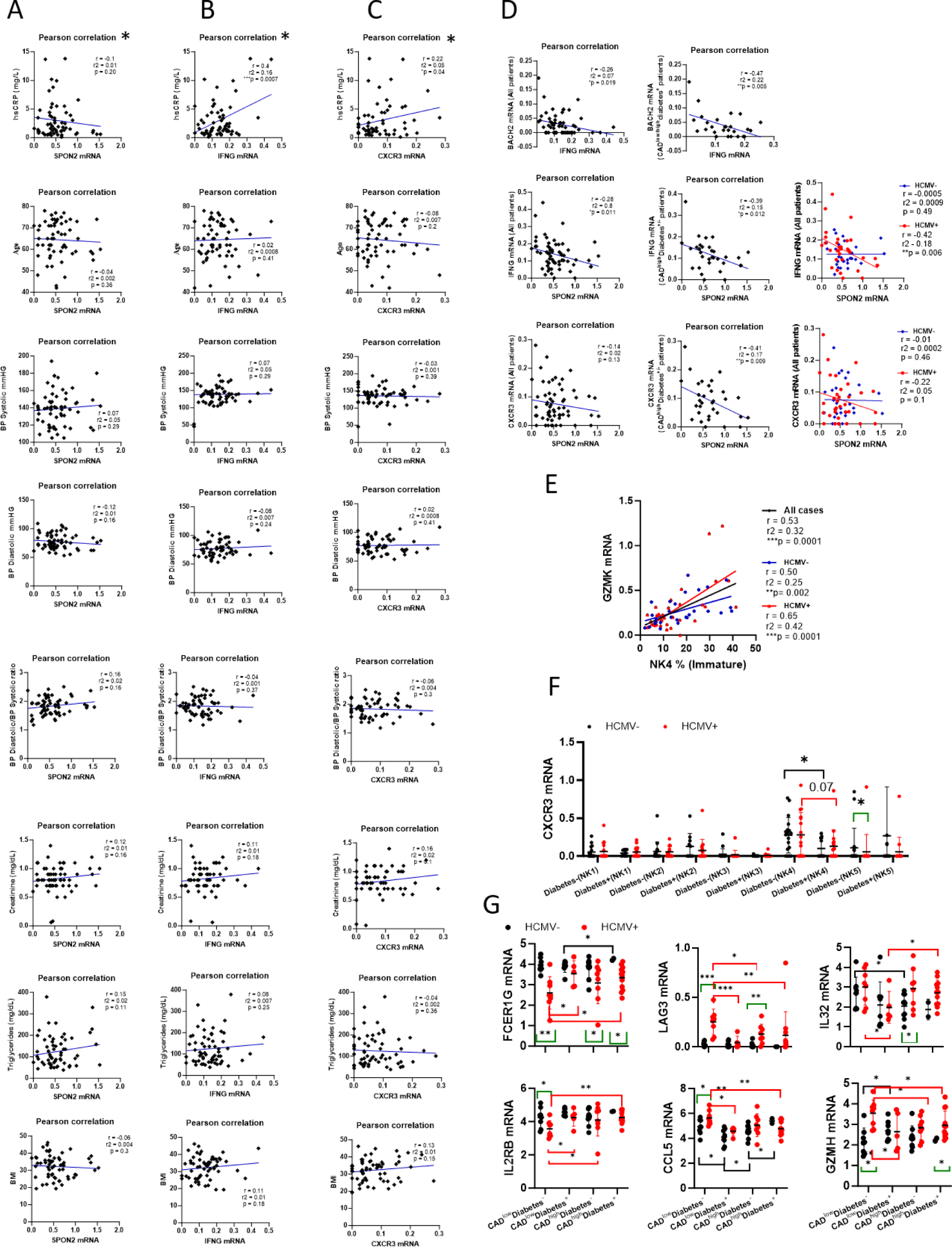
Correlation analysis between **A)** NK cell *SPON2*, **B)** NK cell *IFNG*, or **<u>C)</u>** NK cell *CXCR3* mRNA expression relative to (upper to lower) hsCRP (mg/L), patients’ age, BP Systolic (mmHG), or PB Diastolic (mmGH), BP Systolic/ PB Diastolic ratio, Creatinine (mg/dL), Triglycerides (mg/dL), or BMI values. *To avoid misinterpretation of the data, one patient outlier (hsCRP (mg/L) = 150) was removed from the analysis. **D)** Correlation analysis between; **Upper panels:** NK cell *IFNG* vs. NK cell *BACH2*; left: all cases, right diabetes^+^ cases. **Middle panels:** NK cell *SPON2* vs. NK cell *IFNG*; left: all cases, middle: CAD^high^ cases, right: HCMV^+^ or HCMV^−^ cases. **Lower panels:** NK cell *SPON2* vs. NK cell *CXCR3*; left: all cases, middle: CAD^high^ cases, right: HCMV^+^ or HCMV^−^ cases. **E)** Correlation analysis between cluster NK4 (immature NK cells) proportions relative to NK cell GZMK mRNA expression: All cases (black), HCMV^+^ cases (red), HCMV^−^ cases (blue). **F)** Expression of CXCR3 mRNA in each NK cell cluster (NK1-NK5) in diabetes^−^ vs. diabetes^+^ patients and HCMV^−^ (black) or HCMV^+^ (red) patients. **G)** NK cell mRNA expression of adaptive NK cell signature genes (*FCER1G*, *LAG3*, *IL32*, *IL2RB*, *CCL5*, *GZMH*) between the patients’ groups and HCMV serostatus (black HCMV^−^, red: HCMV^+^). statistical analysis: person correlation (one-tail) or mean+/− S.D, Mann, Whitney test, one-tail (* p <0.05, ** p <0.01, ***p <0.001, green bars: HCMV^−^ vs. HCMV^+^, Black bars: between HCMV^−^ patients’ groups, red bars: between HCMV^+^ patients’ groups).

**Supplementary Figure 2:**
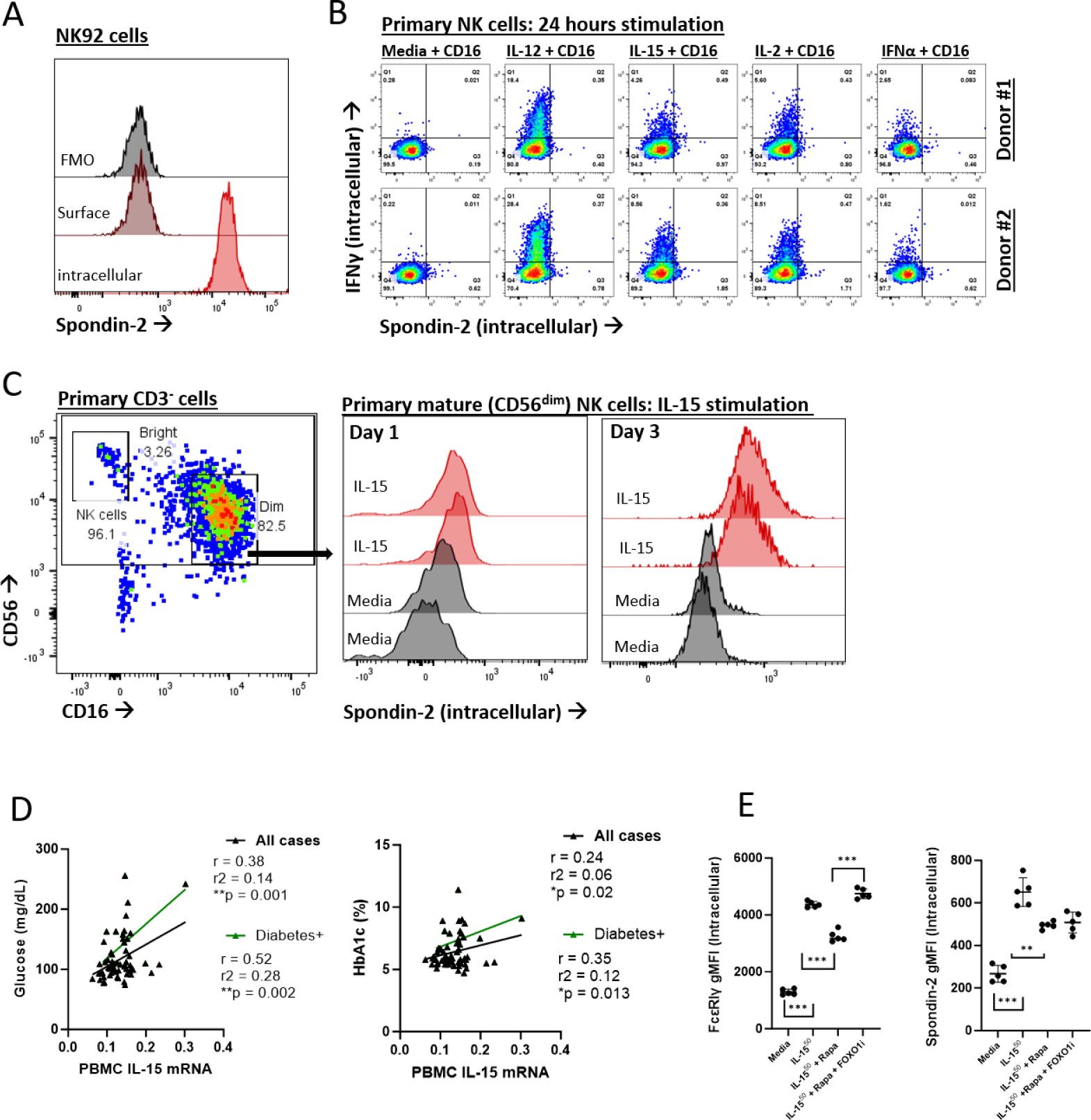
A) Histograms of Spondin-2 intracellular expression (light red) relative to Spon-din-2 surface expression (dark red), relative to FMO control, in NK92 cells. B) Dot plots of intracellular expression of Spondin-2 (X-axis) vs. IFNγ (Y-axis) in isolated human primary NK cells following 24 hours of CD16 stimulation with media without cytokines or with IL-12 (1 ng/ml), IL-2 (300 U/ml), IL-15 (50 ng/ml), or IFNα (50 ng/ml) in two donors. C) Histograms of intracellular expression of Spondin-2 in mature CD56^dim^CD16^+^ primary NK cells (let dot plot) after 1 day or 3 days of stimulation with IL-15 (50 ng/ml) relative to media without cytokines. D) Pearson correlation (one-tail) between PBMC IL-15 mRNA expression and glucose (mg/dL) or HbA1C % in all patients (black line) or diabetes+ patients (green). E) Upregulation of (left) FcɛRIγ or (right) Spondin-2 expression in purified primary NK cells stimulated for 6 days with IL-15 (50 ng/ml) with or without rapamycin (RAPA, 10 nM) or FOXO1 inhibitor (50 nM) relative to media without cytokines.

**Supplementary Figure 3:**
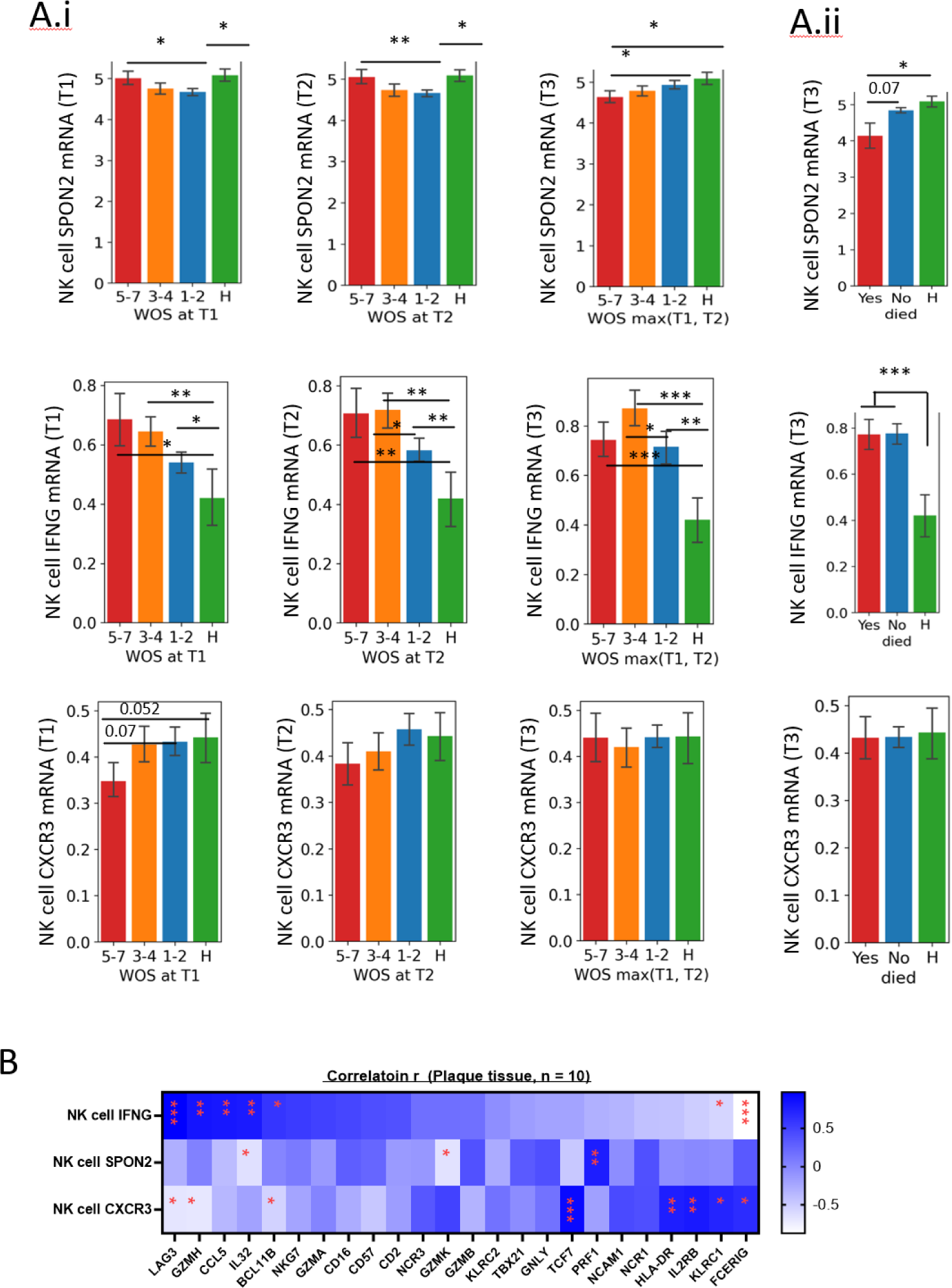
A) i; NK cell *SPON2* (upper panels), *IFNG* (middle panels), or *CXCR3* (lower panels) mRNA expression at COVID-19 cohort T1 = diagnosis, T2 = follow-up, one week after diagnosis, and T3 = long-term follow-up, 2-3 months after initial diagnosis, and relative to COVID-19 disease severity (WOS score: H = healthy, 1-2 = mild, 3-4 = moderate, 5-7 = severe). ii; NK cell *SPON2*, *IFNG*, or *CXCR3* expression at T3 relative to patient’s death. Mean+/− S.D, Mann Whitney test, one-tail (* p <0.05). **B)** Heatmap showing person correlation (one-tail) r values between NK cell *IFNG, SPON2, or CXCR3* mRNA expression relative to the indicated genes in the carotid and femoral plaques (n = 10). *p < 0.05, **p < 0.01, ***p < 0.001

